# HaloUMI: Physics-informed analysis of inhibition halo assays

**DOI:** 10.64898/2026.08.08.743694

**Authors:** Alex Pembery, Hatwan Nadir, Chris MacDonald, Mark C. Leake

## Abstract

Quantification of microbial growth inhibition is central to assays from antibiotic susceptibility to antifungal sensitivity, yet existing approaches struggle with irregular inhibition zones and variation in microbial lawn density. Here, we present Halo Unbiased Measurement of growth Inhibition (HaloUMI), an open-source Python graphical interface for automated, high-throughput analysis of lawn-based microbial assays. HaloUMI integrates robust image processing with physics-informed modelling to quantify inhibition zones without assuming circular geometry, enabling analysis of uniform and irregular halo phenotypes. Using diffusion- and growth-based physical modelling, HaloUMI experimentally validates a correction for variation in microbial lawn density, a major source of assay variability that can confound quantitative comparison of inhibition phenotypes. Validation using simulations and yeast killer-toxin assays demonstrates precise, reproducible measurement across diverse conditions. HaloUMI is applicable to multiple assay formats, including microbial interaction, mating, and conventional disc-diffusion assays, providing an accessible and generalisable framework for quantitative analysis of microbial growth inhibition.

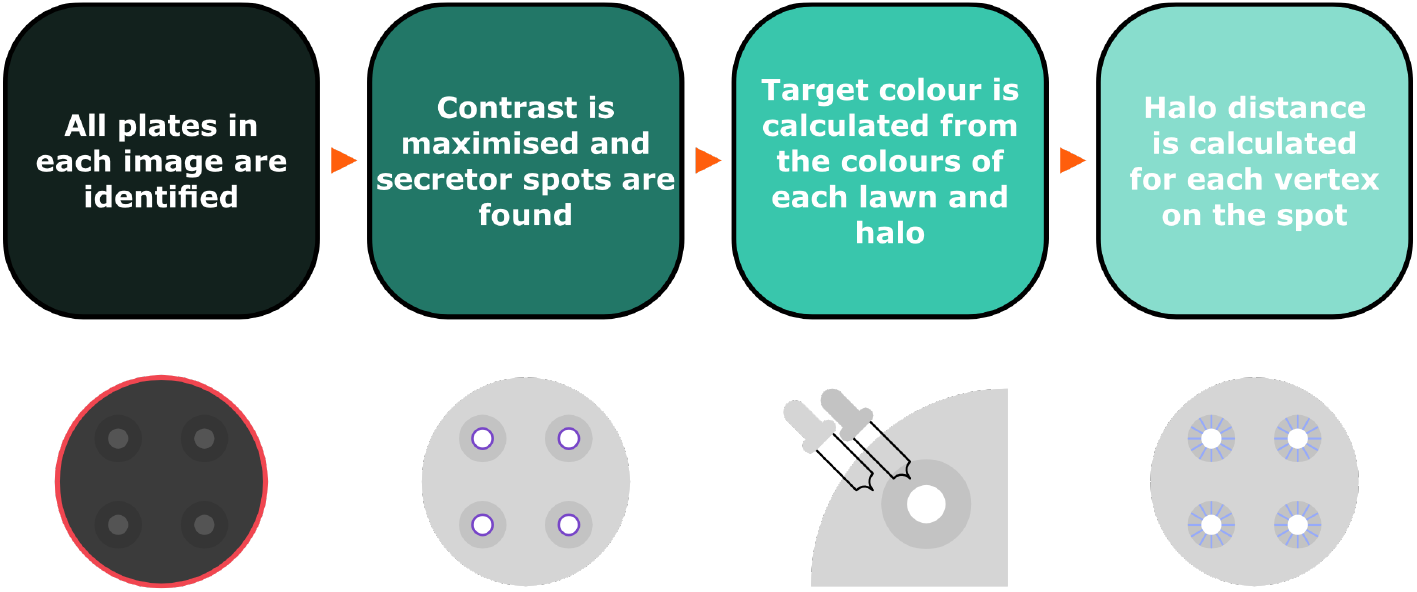

## 1 Introduction

Zone of inhibition assays are typically used to determine the susceptibility of microorganisms to various antimicrobial agents. By measuring the absence of cellular growth on an agar media plate around a disc of filter paper impregnated with an antimicrobial, the efficacy of different compounds can be assessed [1]. This method was standardized by Kirby and Bauer in the 1960s and has since become a reliable system, accessible, and ubiquitous tool. These ‘halo’ assays are applied across a wide variety of biological contexts, from antibiotic efficacy, to showcasing differences between bacteria and archaea [2], screening of enzymatic activity [3], and testing clinical isolates of pathogenic fungi to antifungal compounds [4]. Beyond these valuable contributions, halo assays can be insightful introductory assays to students, and an economical method for elucidating phenotypic differences; invaluable for antimicrobial resistance research in low- and middle-income countries [5].

The annular radius, or halo, around the chosen reagent is defined by the critical concentration; approximately equal to the minimum inhibitory concentration obtained in broth dilution susceptibility tests [1]. The spot of reagent is often well-defined by the filter paper disc, or in the case of a well-diffusion assay, a punched-out cylindrical hole in the agar in which the reagent can then be dispensed [6]. Both of these methods are characterized by the uniformity of the region of antimicrobial, and correspondingly result in a circular halo. As such, they are typically measured with a ruler or calliper measurement [7], as is common practice throughout school and university education [8]. However, the time-consuming nature of repeated manual measurements and the influence of human-error [9] has inspired multiple different approaches to tackle this problem. Some systems rely on the simplicity of the assay by utilising the circular nature of the spots [9, 10], allowing for simple detection and analysis of the halos using methods like cv.HoughCircles [11]. Others have implemented neural networks [12] and more complex methods such as laser speckling [13, 14], in order to increase throughput and accuracy. Whilst zone of inhibition assays are typically used for observing antimicrobial activity in bacteria, they can also be used to study the biological processes in the budding yeast *Saccharomyces cerevisiae*. For example, a halo-based method is routinely used to define the mating type of different yeast species, such as testing pheromone responsiveness by using filter paper discs of different *α* factor concentrations on lawns of MATa strains [15]. Another example is measuring the effects of secreted toxins on other yeast cells. One example includes the effects of the yeast killer toxin K28 exposed to yeast mutants. K28 is secreted by *Saccharomyces cerevisiae* cells hosting ScV-M28 and its L-A helper virus [16, 17]. The mechanism behind the toxicity of K28 is sensitive to pH due to theorised disulfide-bond rearrangements in order to release the toxic *α*-subunit [18]. The requirement for mildly acidic conditions and preference for rich media [19] mean that the use of a paper disc is suboptimal. Whilst the well-diffusion assay is an attractive solution, the high-degree of sterility required for creating the wells is paramount and adds complexities unideal for high-throughput screens.

Instead, it is shown that halo assays can simply be performed by pipetting a spot of toxin secretor cells that have been cultured separately onto the lawn of cells [20], thereby being both as economical in time and resources as the other two aforementioned protocols. However, the perfect circles present in those methods are now absent due to the myriad physical factors that govern how a small volume interacts with the agar plate. Non-uniformly circular spots thereby create difficulties when measuring the diameter, which can be illustrated in the simple example of an oval, wherein its dimensions are described by two axes, not a singular diameter that can be measured. As the complexity of these spots increases, so does the number of measurements required to garner a precise picture of the sensitivity of a lawn of cells to the toxin. In the past, we have done this through multiple spots of secretor cells per plate, and taking at least four measurements per spot; one for each cardinal direction [19]. Thereby for just one lawn of cells, 50 individual readings are taken using ImageJ before being used to create an average. To observe the effects of between different gene families or functions, this could require upwards of 100 different deletion mutants; presenting a vast amount of manual work, in which human error can easily creep in. This is exemplified in the Carroll *et al*. K28 screen [21], wherein the limited resolution and low degree of quantitativeness resulted in false negatives [19]. This error is subsequently heightened by an unconscious confirmation bias, or if measurements are collated from multiple people, as might be the case for a workshop or practical, since it can be difficult to determine the where the halo edge lies.

Consequently, the spotted version of the halo assay is highly inclined toward an automated system, yet the “complex simplicity” is not trivial to solve. The closest solution to the problem is addressed by Costa *et al*. [22] wherein they propose an automated system that finds the colour at which an averaged vector transitions from the dead cell region to that of the lawn, and applies this to four arcs of a circle to gain an accurate picture of the halo. Although this elegant system still assumes for a perfect halo circle, it has inspired our efforts to segment irregular shapes, from the image preprocessing to the flexibility in assessing diameters, both of the halo *and* the filter paper disc. We also note this alternative system is not publicly accessible, and the number of measurements generated are insufficient to accurately determine K28 sensitivity in our assays. We were similarly inspired by Gerstein *et al*.’s *diskImageR* [23] which clearly quantifies measurements for single disc diffusion assays using radial measurement lines. They successfully show how this can be used to distinguish between drug susceptibility as well as further exciting developments such as inferring minimum inhibitory concentrations. This, however, is again done for perfect circles, lacks the capability for high-throughput analysis, and does not correct for any experimental variation. Consequently, we seek to create a similar pipeline whilst also addressing some additional parameters and championing accessibility as a priority.

Therefore, we present here that of HaloUMI (Halo Unbiased Measurement of growth Inhibition); a Python-based system that performs hundreds of readings per plate, accurately calculating the halo radii for dozens of points around a spot.

## 2 Results

### 2.1 Analysis Pipeline of Code

#### 2.1.1 Contour Filtering and Identification

To facilitate high-throughput analysis, an automated method was required to isolate individual culture plates from the original TIFF file (Figure 1a) that typically have between 1 to 6 plates. The raw image data underwent initial formatting and normalisation to 8-bit depth before a temporary resizing to optimise efficiency when identifying each plate with a Hough Circle Transform (Figure 1b) [24]. This method is constrained by requiring plate centres to be a set number of pixels apart, and a small window of freedom for the sizes of the plates themselves outlining the geometric parameters.

**Figure 1:**
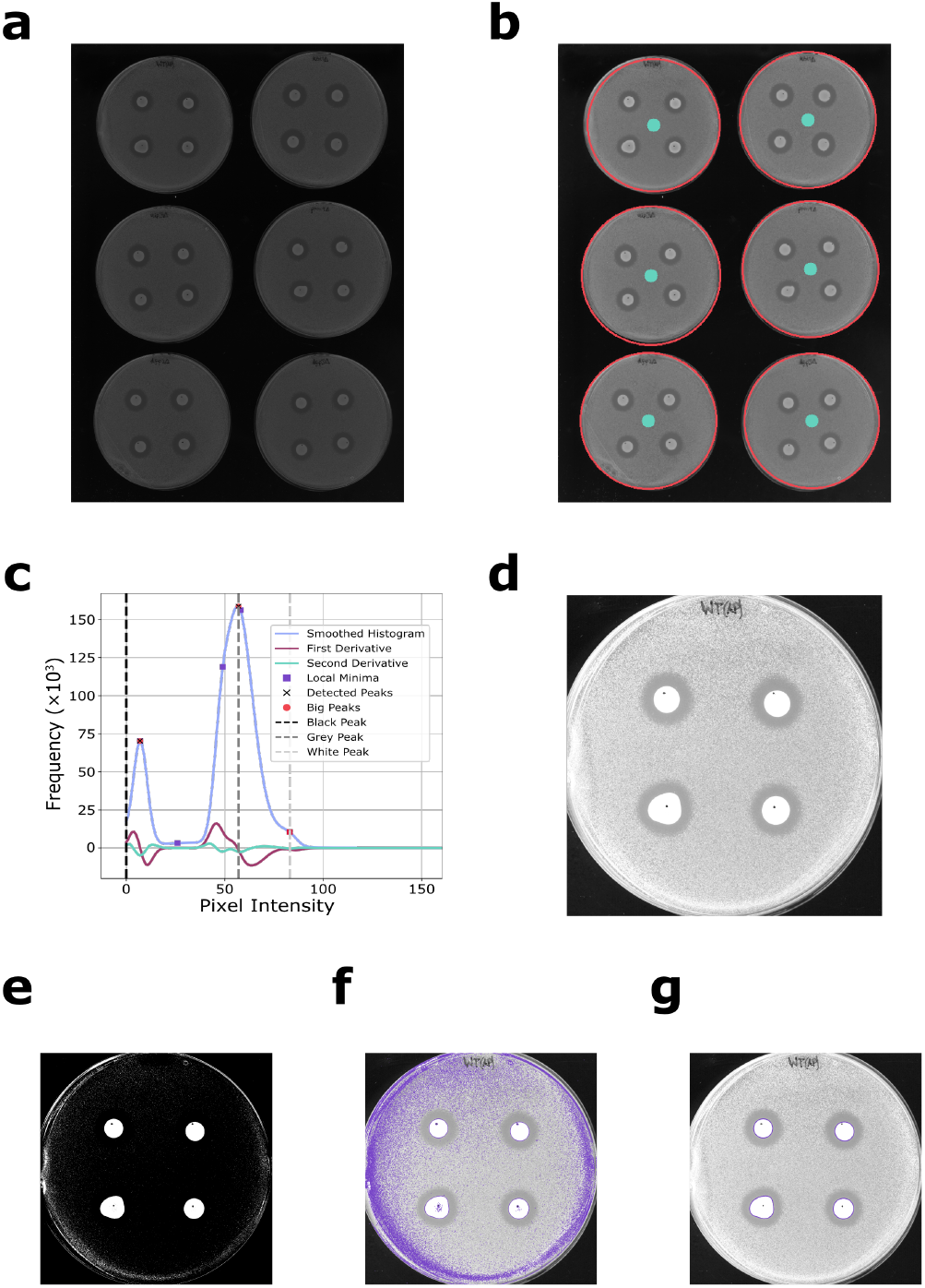
Stepwise breakdown of detection of select non-uniform contours. **a)** Original TIFF file taken by scanning in plates on a charge-coupled device scanner. Files are 16-bit greyscale, 600dpi and 5100px x 7019px. **b)** A 2x contrast is applied, and the plates are identified using the open computer vision library, the centres and radii are outlined in teal and red, respectively. **c)** SciPy was used to create a smoothed histogram created using 1D Gaussian smoothing. Relevant maxima were detected using SciPy’s find_peaks. **d)** Setting a maximum brightness to the average of the grey and white peaks is used to generate a highly contrasted plate without losing important experimental detail. **e)** Binary image for detection of K28 secretor cells, wherein any pixels with intensities below the threshold are black, and white otherwise. **f)** All detected contours found using Opencv.The increasing frequency of contours towards the edge of the plate is indicative of the non-uniformity of the lawn, creating gaps that the algorithm interprets as miniscule contours. **g)** Filtered contours resulting from a thorough screening using the height, width, ‘radius’, and area

Artifacts like minor surface defects often result in locations of increased brightness, thereby skewing the image to appear darker. This is overcome by implementing histogram equalization to create a repeatable method for contrasting (Figure 1c). Each image contains three distinct regions: a black background, a grey lawn, and the white spots of cells or discs. The black and grey regions dominate and as such are mostly trivial to identify, after smoothing and locating with the gaussian_filter1d and find_peaks SciPy [25] functions respectively. Depending on the image, this may typically produce one or two peaks, depending on if the black peak is undetected. This may occur if the peak is centred about 0, instead of producing a symmetrical peak centred in the low tens of brightness. As such, the grey peak must be assigned either as the first or last peak exceeding a set frequency, whereas the black peak is always set to zero. A higher contrast may be achieved by replacing the black peak with a value in between zero and the grey peak. However, to consistently manufacture this across a diverse range of scanners and plates would require a further refinement, where it is arguably not necessary since the resulting contrast from this pipeline is already more than satisfactory (Figure 1d). The binary plate (Figure 1e)is used to locate all defined contours using the findContours function from Opencv.This accurately identifies the key spots, albeit whilst also including contours from the plate outline, surface defects, and edge effects (Figure 1f). Consequently, a filtering function is required to isolate the spots from the noise, the parameters of which can be defined by the anticipated morphologies of the spots themselves, namely the expected areas, ‘diameter’, width, and height.

The area is calculated by approximating the spot as a polygon using cv.arcLength and cv.approxPolyDP. A suitably small value for the vertex factor *ϵ* can be set such that the polygon is not an oversimplification yet not too complex that computing times are suboptimal. Valid areas must satisfy the inequality of being between 0.2% and 5% of the area of the image. This gives significant flexibility necessary for variations in size attribute to the use of pipetted drops.

These contours must be further filtered to remove those that have valid areas yet unsuitable morphologies (i.e. a circle of radius 4cm has approximately the same area as a rectangle of side lengths 5cm and 10cm). In brief, the spots must resemble a circle, typically achieved with a roundness score using calculations involving the perimeter, though this can become inaccurate when considering the coastal paradox. Therefore, the ‘diameter’ of the spot is approximated simply as 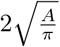, with which it is used in ratiometric inequalities with the width and height of the bounding box of the spot, wherein each ratio should lie between 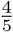 and 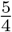 to account for some natural variations. If both of these checks are cleared, then the centres of each of these spots are found and a final check is performed to ensure that the spots are significantly apart from one another (Figure 1g).

#### 2.1.2 Halo Radius Detection and GUI

The method of detection is defined as the point exactly between the ‘deadzone’ (the region of no growth) and the lawn. To implement this, the average colour of *both* of these areas must be calculated. Since there can be variations in intensity across the plate resulting from how the plate was poured or the lawn preparation, it is essential to calculate these colours for each of the spots. This is achieved by drawing averaging contours of the same shape as our secretor spot, but of increasing size (Figure 2a). The deadzone ring must be large enough that it avoids the gradient of the toxin, yet small enough that it can be used for resistant phenotypes. Based on multiple refinements, a multiplier of 1.25 was chosen, which thus also represents the limit of detection. This subsequently explains the choice of ratios lying between 1 ± 0.25.

**Figure 2:**
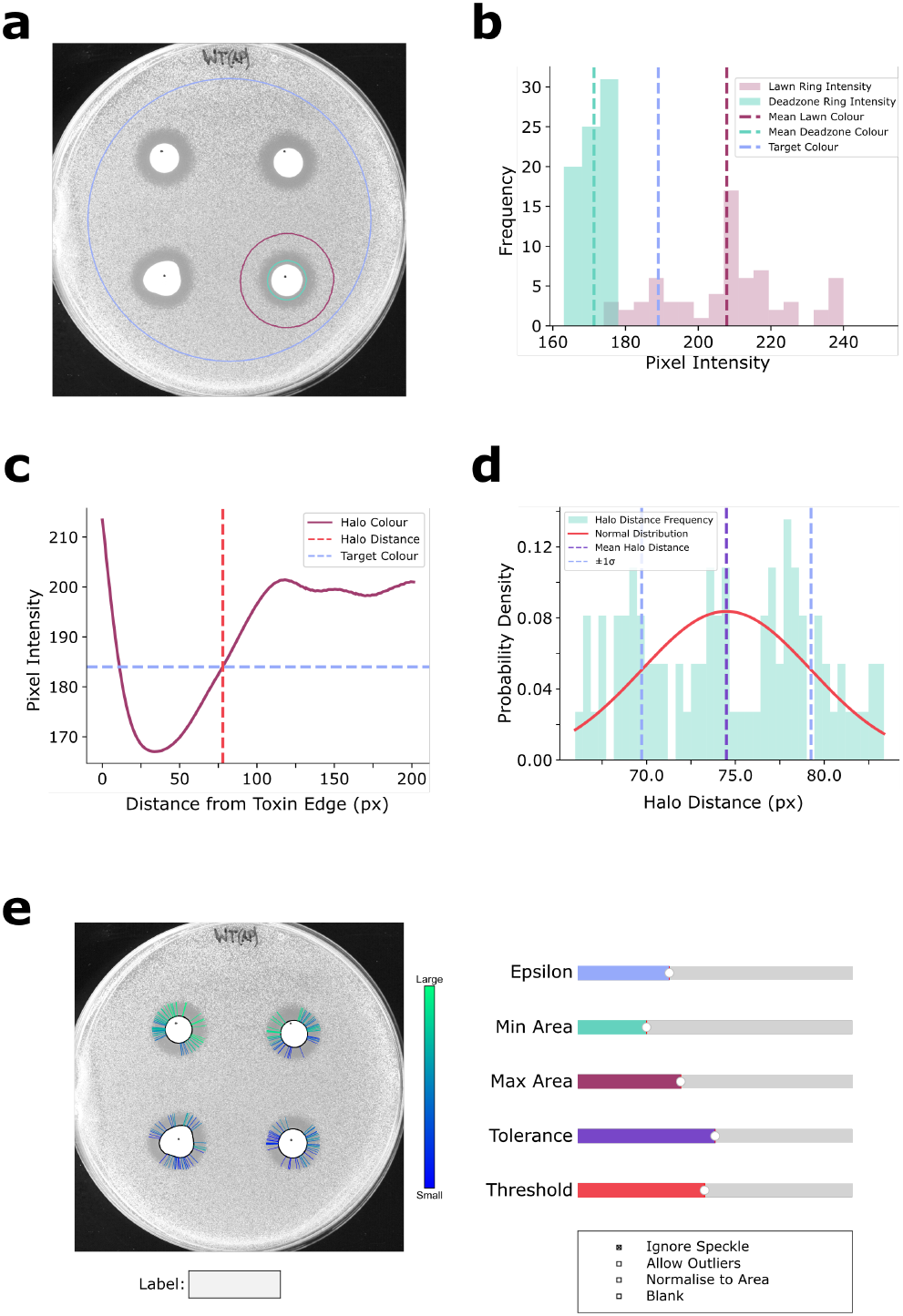
Method of halo detection and corresponding results. **a)** Expanded contours for the calculation of the target colour, with the dead zone and lawn colours calculated from the green and dark red contours respectively. The blue contour represents the 80% plate limit. **b)** Histogram showing the distribution of the deadzone and lawn colours, with the target colour exactly halfway between the two. **c)** Pinpointing of the halo distance, with a line profile for a singular vertex drawn in dark red, showing the exact point it exceeds the target colour. **d)** Gaussian distribution of all halo distances for one plate. **e)** The graphical user interface features five sliders that respectively control the number of vertices on the secretor spot polygon, the minimum area, the maximum area, the allowed ratio for the contour filtering (and its reciprocal), and the threshold value used for contrasting and binarization. Four tick-boxes are also provided for the user to: ignore the filter for a stringent lawn colour, to allow any filtered outliers, to normalise all values per spot to the area, and to visually clear the halo distance lines for an alternate view.

Similarly, the lawn ring must be similarly controlled: it must not exceed the boundaries of the plate but still lie in the deadzone for the most sensitive mutants. Consequently, a multiplier of 3 was chosen. Whilst this may be suitable for perfectly placed spots, this system is designed to be as versatile as possible meaning that spots may be larger than others, or may be closer to the edge than desired. Since most surface defects are found closer to the edge, it is also required that the lawn ring cannot be closer than 80% of the plate radius to the edge. Should this be exceeded, it simply results in a truncated ring for which to calculate the average colour of the lawn.

To ensure halos are distinctly defined, a final check is implemented. The lower bound of the lawn *µ*_*lawn*_ −*σ*_*lawn*_ must be greater than the upper bound of the deadzone *µ*_*deadzone*_ + *σ*_*deadzone*_, and the lawn coefficient of variation 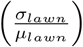 must be less than 0.25. This eliminates spots with noisy lawns such that halos are not artificially inflated by surface defects. However, in rare cases such as when the lawn density is too disperse, this check can sometimes be impossible to clear. To allow the user to determine if this is an issue, a toggle is built into the GUI to allow for a filter bypass.

Regardless, the target colour (Figure 2b) is always calculated as:

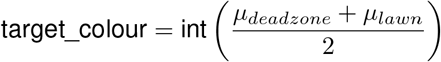

Due to the uneven nature of this phenotype, a new method was required to instead calculate the halo distance from the edge of the secretor to the start of the lawn, from *each* vertex on the secretor spot. This highlights an additional benefit of approximating the spot as a multi-vertexed shape. To ensure consistency, it is imperative to define the angle at which the halo distance is calculated from each vertex. This was initially achieved by defining the halo distance as the shortest possible length from the secretor to the lawn, calculated by considering the angle perpendicular to the surface, since any fluctuations in angle would theoretically result in a greater halo. Whilst this worked for the majority of vertices, it failed when parts of the surface were concave since the angle would point away from the lawn, and result in skewed distances. As such, a less mathematically perfect approach was necessary for the highly variable assay. Consequently, the angle is taken to be the unit vector from the secretor centre and the vertex. This can be thought of as measuring the halo *radius* from the secretor centre, then subtracting the distance it took from the centre to the secretor edge.

A line of 200 points is then drawn from each vertex and the x and y values for each point are calculated using the unit vector. These points are checked with a range of if-statements to ensure their suitability. If all points are deemed to be valid, then the distance and colour of each point can be calculated (Figure 2c). To find the colour at each point, instead of just observing the colour at the pixel, an averaging mask is used to smooth the surroundings, such that any surface defects are avoided. This is because the detection method is looking for the first instance of a pixel that supersedes the target colour, hence any anomalous marks would likely result in a false positive.

To calculate the halo distance, it is first necessitated that the distance must be greater than 25% of the secretor radius, in line with the limit of detection. It then just remains to be worked out the point at which the target colour is reached. This was initially done by observing the first point to have a brightness greater than the target colour. However, since this method is based on discrete points, two distinct populations were forming: one where the target colour was only just cleared, and another where the target colour was significantly cleared, yet the previous point had not. To counter this, a linear fit is applied to the two points either side of the target colour and the distance at which the target colour is met can be interpolated as the halo distance.

As a final filter, a halo distance is only marked as valid when it is within one standard deviation of all other distances from that spot (Figure 2d). Furthermore, the coefficient of variation must again be less than 0.25 to ignore spots that exhibit multiple populations, likely resulting from surface defects or gradients in the lawn density. If this filter is passed, then the halo is drawn on the image, contributing to the annotated copy for presentation purposes.

Despite allowing for leeway in many of the variables through the use of inequalities, the highly diverse nature of the experiment means that these may not catch all possible cases. Extensive testing showed that this is notably the case when considering the minimum and maximum areas of the secretor spots and the calculation for target colour requiring a small standard deviation. Consequently, a simple GUI is implemented to allow the user to easily adjust these parameters, as well as the number of lines drawn, aspect ratio tolerance, and whether to include the outliers removed in the final step (Figure 2e). It must be noted that when using the GUI that for the experiments to be repeatable, similar settings should be used throughout, especially when modifying the Epsilon sliders or allowing outliers, since this has a global effect, instead of just missing out some of the secretor spots. However, this caveat is minor is comparison to having an automated system that would have to be constantly re-run and modified each time.

### 2.2 Simulated Data

Simulated examples were generated, using a vector graphics editor, to assess the robustness of HaloUMI and demonstrate its capabilities. Example shapes can also be easily formulated to highlight the morphological filter (Figure 3a). The simplest variation is large halos with clearly defined spots and no surface defects (Figure 3bi). As expected, highly consistent results with a coefficient of variation of just 0.05% and a percentage error of just 0.006% are observed. The small degree of outliers likely stems from the quantised pixels and represent a tiny error window of the pipeline. If these are included, then the coefficient of variation is still only 0.3%. The next variation is a plate with the smallest possible halo that the code can theoretically detect (Figure 3bii). The percentage error increases to 2.3% in this case, but this is likely significantly less than would be the case for manual measurements.

**Figure 3:**
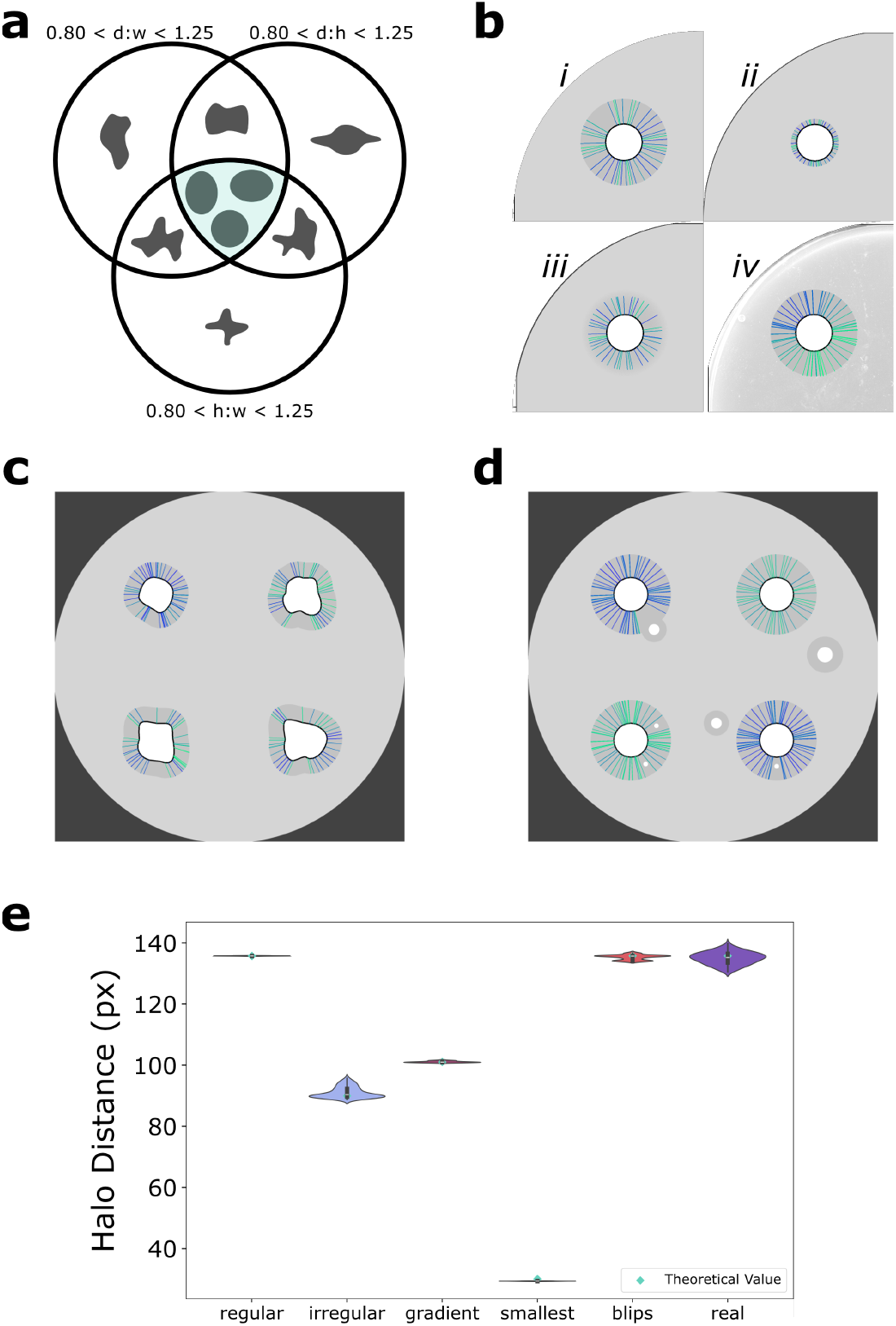
Simulations for exploring code functions. **a)** Venn-diagram to represent the different ratiometric equations of diameter, height, and width, and that the code only allows spots that satisfy all three inequalities (in green) **b)** i) typical-sized halo, ii) smallest detectable halo, iii) halo with a gradually fading edge, iv) halo with a real background taken from a scan of an empty plate **c)** Simulated spots with extreme cases of unevenness that still successfully meet the morphological criteria **d)** Simulated plate with ‘blips’ of secretor cells, which the code is successfully able to detect and avoid **e)** Violin plot representing the theoretical values of each plot as determined by the Affinity software and the actual values calculated by feeding the simulations through HaloUMI

Since these are biological assays, the lack of binary must also be considered. We find that there is often something of a gradient between the deadzone and the lawn, stemming from cells that lie at the threshold between lethal and sub-lethal concentrations of toxin. Cells at this threshold region present an issue in marking the end of a halo. In Figure 3biii, it perhaps looks as though the code has underestimated the boundary. Whilst this is an appropriate assumption, creating an algorithm that is consistent in its detection with a vastly more complex biological system, it much trickier to implement. As such, so long as it is repeatable, the outputted data are more trustable than manual measurements, which is why simple definition of the halo distance (the length from the edge of the spot to colour exactly halfway between that of the deadzone and the lawn) suffices. Whilst it would be trivial to define it as any fraction along the line, having a consistent variable again helps with repeatability. Further research into the biology itself could elucidate the optimal point of crossover and better reflect the complexity of the environment (Figure 3biv).

The previous examples all have the benefit of being perfectly circular in morphology; we next addressed asymmetry and amorphous regions (Figure 3c). Mathematically and biologically, we cannot just consider halos to be scaled versions of the spot since they are not radially symmetric. Instead, the halo is better described as the Minkowski sum of the spot since at all points on the perimeter, there will be cells that secrete toxin. This does make it more challenging to consider the normal to each point on the surface, yet with the more forgiving and round examples of the spots in our experiment, it is deemed to be an appropriate approximation.

Figure 3d highlights the necessity of the removal of outliers. Although the averaging mask helps to smooth any bright surface defects, if a mark is large enough, then this cannot simply be smoothed out. Such examples may occur when a small amount of secretor cells from the tip of the pipette is accidentally dropped onto the surface. Since the halo distance is determined at the point at which the target colour is exceeded, even just one surface defect could cause an early spike. Previous iterations considered the requirement that to satisfy being designated as the lawn, the colour must plateau for a certain number of pixels. However, the complexity that this would require for repeatable measurements made it challenging and ineffective to implement. As such, the code simply allows for these smaller and larger halos to be detected but they are eliminated in the final valid halo filter.

The described examples are just some of the possible variations that we may encounter, and each addresses the potential issues as stand-alone (Figure 3e), when in reality they are often compounded with each other. We therefore must upgrade our simulations to working with well-described, real data.

### 2.3 Real Examples

By running controls of wild type cells and cells lacking a K28 defence factor (*ktd1*Δ[26]), we can assess the importance of this gene on killer toxin defence through analysing the difference in halo size. By normalising to WT each time, we can observe that the difference between WT and *ktd1*Δ is a factor of approximately 1.3. This is a significant difference and is indicative of a protein whose function is directly related to defence, as is expected for Ktd1 (killer toxin defence). Whereas, the difference between wild-type and an unrelated gene may be in in the region of 0.9 - 1.1, suggesting it is akin to wild-type in terms of defence. By using HaloUMI, we have been able to screen multiple groups of proteins, and assess the importance of any strains that are significantly sensitive or resistant.

Through optimising the code, it has also been possible to further optimise the assay. It is well-known that the more dense a lawn is, the smaller the size of the halo. This is a combination of two factors: 1) a denser lawn will reach a critical concentration at which the halo is visible faster [27]; 2) the minimal inhibitory concentration (or critical concentration) increases significantly as the number of organisms is increased (inoculum effect) [28]. Therefore, since it is desirable to have a large dynamic range between resistant and sensitive phenotypes, large halos are most desirable, thereby requiring a sufficiently diffuse lawn.

In Figure 4a it is clear to see the size difference between wild-type and *ktd1*Δ and the effect of the lawn. This highlights just one application of this assay and the resulting data, as well as the flexibility and wide-ranging function HaloUMI can have. Figure 4b demonstrates just some uses of HaloUMI.

**Figure 4:**
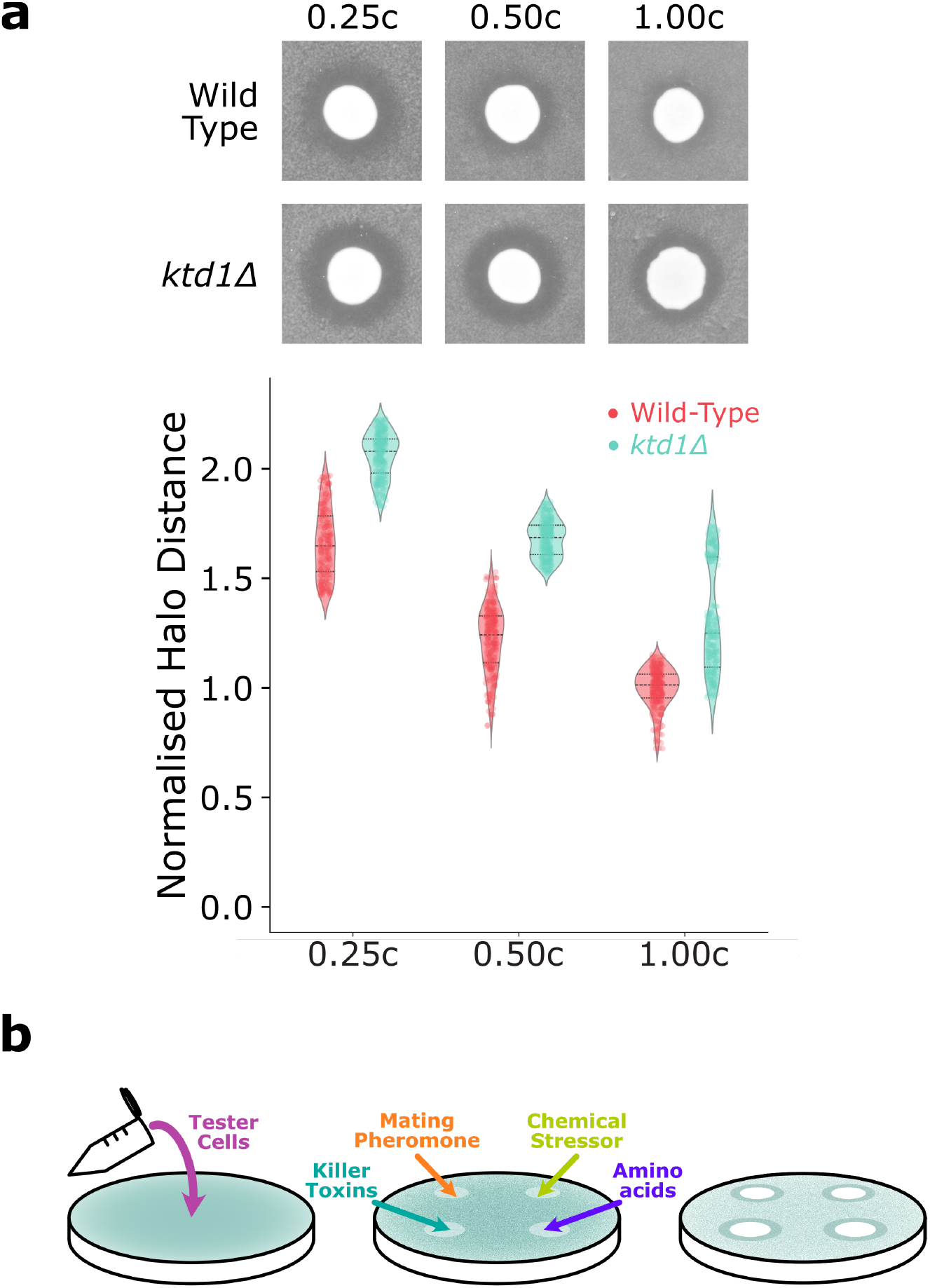
Simulations for exploring code functions. **a)** Comparison between wild type and cells without killer toxin defence factor (*ktd1*Δ) **b)** Method for carrying out halo assay, with different examples for the spots shown that would result in similar yet distinct phenotypes

Not only can spots be pipetted, but they can also be made more traditionally through either wells or discs. These myriad use cases are ideal for a GUI, allowing the user complete control over their results. To improve on this further, a point-and-click interface is currently being developed.

## 3 Discussion

In this study, we document a high throughput analysis pipeline for the assessment of microbial growth inhibition assays. Using the sensitivity of *Saccharomyces cerevisiae* to virally encoded killer toxin K28, regions of lawn cell death were used for optimisation. This system improves precision, throughput, whilst reducing human error. The analysis package can account for abnormal morphologies of biological samples as well as providing a novel normalisation calculation for variations in lawn density. This package offers an accessible graphical user interface that has broad utility for related analyses.

Whilst great care is taken to normalise the lawns to a set concentration each time to ensure consistent and reproducible data, small fluctuations can greatly impact the actual concentration of the lawn. As such, if a strain was for example to be too diffuse, it might result in a relatively overinflated halo. Previously, this would be hard to detect and equally hard to adjust. To correct for this, we must apply a physical model.

To approximate the concentrations of the lawns and secretor cells, a point-source diffusion model is implemented, as shown by Gefen *et al*. [27]:

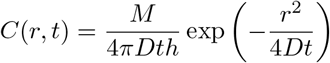

*Where C is the concentration of antibiotics as a function of time (t) and distance (r) from the source (centre of disk), assuming isotropic diffusion in a two-dimensional plate. D is the diffusion coefficient, M is the initial antibiotic amount in the disk and h is the height of the agar in the plate*

This equation is derived from Fick’s 1D diffusion equation [29]. In reality, this relationship is far more intricate since we do not have a point source of an antimicrobial, but rather an amorphous disc of secretor cells. One could argue that we must also account for the coffee-ring effect, as well as the fact that there is less competition for cells on the periphery of the disc compared to those confined in the centre. However, since the area of this disc is orders of magnitude smaller than the whole of the dish, it can be argued that it is still a suitable approximation to make.

Consequently, we can extract some expected relationships from this formula and use HaloUMI to assess its accuracy. Firstly, to use this equation in its simpler form, we endeavour to keep the height of the agar consistent. Additionally, since we typically use halo assays to assess the lawns as opposed to the killer toxins themselves, we can treat the diffusion coefficient as a constant. Now, we can clearly see that if we were to increase the amount of the secretor cells we have, then we would expect to see an exponential increase with respect to halo distance squared. This makes intuitive sense since more killer cells results in more toxin being secreted before a lawn can form sufficiently. However, the non-linearity does show that changes when concentrations are smaller have a much greater effect on the halo distance, perhaps inadvertently representing how the cells will eventually pile up and deplete each other of nutrients, thereby negatively impacting their killing ability.

A further takeaway from the theoretical equation can be made regarding the impact of the concentration not of the secretors but of the lawn of cells being tested. For this we must consider what the halo actually represents. The zone of inhibition represents the interplay between the diffusion rate of the antibiotic, the growth rate of the bacteria, and the minimum inhibitory concentration. As aforementioned, for us this simplifies down to the balance between the growth rate of the yeast, and the MIC of each strain. We can therefore define the halo as the point where the concentration of the toxin drops below the MIC, at the time at which the yeast have grown sufficiently for a halo to be visible [27]. To relate this to the concentration of the lawn, we must consider how long it takes yeast to grow to the visible concentration, *c*_*vis*_. If we assume that yeast are growing in log phase, then the expression for *c*_*vis*_ is simply given by:

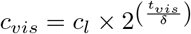

where *c*_*l*_ is the initial concentration of the lawn, and *δ* is the time per division.

By rearranging for *t*_*vis*_, we see that:

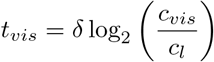

Before appreciating how this relates to halo size, it is important to address a few approximations. We assume that yeast are growing in log phase, however there will be a period of time wherein the cells are initially in lag phase. Additionally, denser lawns will enter stationary phase earlier on. Despite this, since the critical window of toxin effect occurs in the earlier hours, this is largely mitigated against. Additionally, since the lawn is left to dry for around 40 minutes, this allows for some added time to allow for reaching log phase before the secretor cells are added. As such, we can treat the start point for both *t*_*vis*_ and *t* as the same time-point, removing the need for an additional variable.

By rearranging the equation for *r*^2^, we can see how the halo distance varies with time:

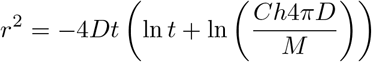

To simplify this, at high values of *t*, the natural log is approximately constant. Therefore, each of the natural terms inside the bracket can be treated as constants. As such:

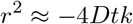

Therefore:

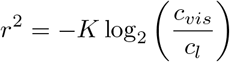

*where K* = 4*Dkδ*

Since in most cases the lawn concentration is aimed to be consistent across experiments, this too can be treated as a constant. Consequently, we find a linear relationship of the form *y* = *mx* + *c*:

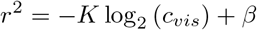

*where β* = *K* log_2_ (*c*_*l*_)

This does still replicate the negative correlation we expect, in that generally, halo distance gets larger as visible lawn concentration gets less dense. This therefore allows us to improve the halo assay further by accounting for the concentration of a lawn. By assuming that the standard deviation of pixel intensities around each spot is a suitable metric for the visible lawn concentration (i.e. as *c*_*vis*_ increases, *σ*_*vis*_ decreases), we obtain:

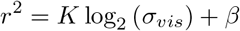

*K* can be calculated by creating a reference set of plates with deliberately modified lawn concentrations and calculating the gradient of *r*^2^ against log_2_ (*σ*_*vis*_). Corrected values for *r*^2^ are then obtained using *K*, relative to wild-type:

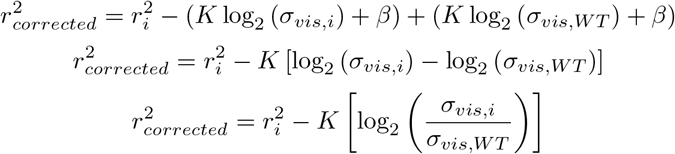

Therefore:

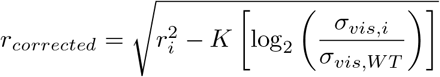

To validate this equation, we can prepare a 4x4 matrix of plates with exponentially increasing lawn and toxin concentrations (L1-L4 (0.0125, 0.0250, 0.0500, 0.1000) and T1-T4 (1.25, 2.5, 5, 10)) (Figure 5a). If we plot *r*^2^ against log_2_ (*σ*_*vis,i*_), the linear relationship is found, from which we can calculate the gradient (Figure 5b). For the purposes of demonstration, let us now assume that our values for halo distance were all performed with the same OD of lawn and we need to correct for any deviations. Using the gradients obtained for each strength of toxin concentration, the *r*_*corrected*_ equation can be applied, essentially levelling the previous positive gradient, to one that is flat (Figure 5c). Whilst this example is incidental, now that values for *K* have been obtained for different toxin concentrations, variations in lawn density can now be accounted for across a multitude of experiments. This will be very valuable in ensuring phenotypes are reported correctly, and that no false positives/negatives are shown.

**Figure 5:**
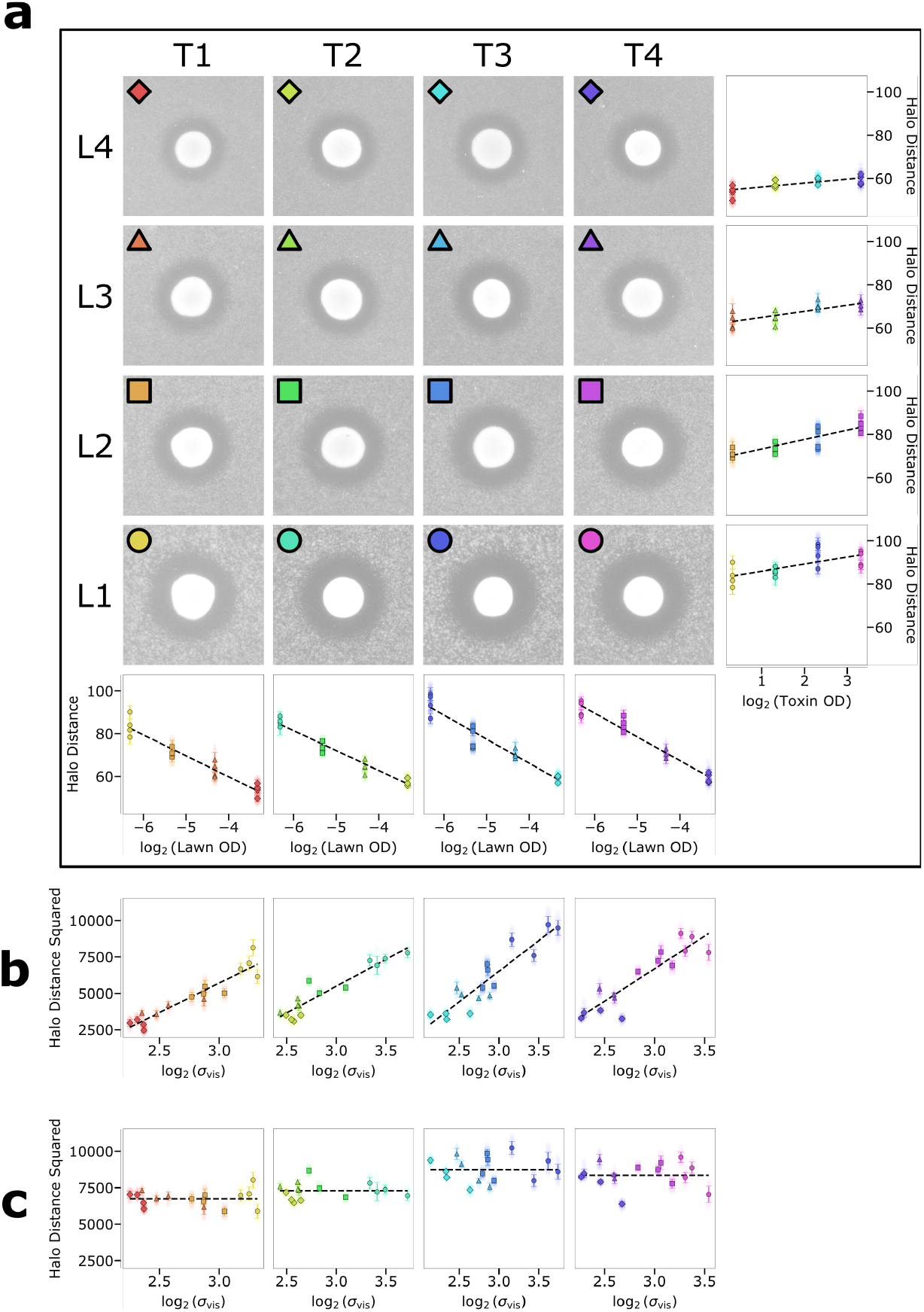
Correcting for variations in lawn density. **a)** Comparisons between different lawn and toxin concentrations (OD of culture prior to plating), where each doubles between subsequent values: L1-L4 (0.0125, 0.0250, 0.0500, 0.1000) and T1-T4 (1.25, 2.5, 5, 10). A snapshot of a typical halo is shown for each, and the collated results from all four spots from each plate is shown in the final column/row. **b)** The square of the halo distance is plotted against the base 2 log of the standard deviation of the lawn ring. **c)** Repeat of plot B but with corrected halo distances, as exemplified by the horizontal gradient.

Another area that can be improved upon is that of the size of the spots. We have found that the hydrophilicity of the agar can vary between experiments, resulting from the age of the plates or autoclaving duration. Consequently, we report that the area of the spot itself is affected by this, and thereby the resulting halo. Consider a hydrophilic surface in which a number of cells in suspension are deposited. The liquid will spread farther and thus the secretor cells are disperse, meaning that fewer cells are on the periphery. If we instead consider a hydrophobic surface, these cells will be more clustered together, and effectively more concentrated. Even accounting for the coffee-ring effect, the amount of cells that contribute toxin to its outer environs is likely much greater and would result in a larger halo. As such, to improve our automated model yet further, we should consider the effect that this may have, particularly when trying to normalise across many days of experiments.

To incorporate this into our model, we can extend our equation and corresponding proportionality relationships. The initial formula treats the secretor spot as a point-source, yet for an upgraded model, we can incorporate the coffee-ring effect and have a system of many point sources, integrated around infinitesimally small sections of a ring. This is analogous to the system for heat conduction for an instantaneous ring source at *t* = 0 of strength *Q′* = 2*πr′Q* and radius *r′* in the plane *z′* = 0, as described by Carslaw and Jaeger [30]:

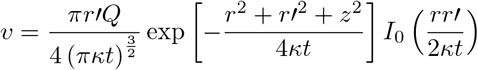

We can simply translate this into our semi-3D system, making sure to note that due to the competition in the centre of the spot, we only consider where *r > r′*:

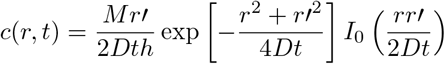

Comparing this to the point-source equation, it is clear to see the increase in complexity:

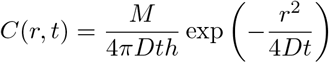

More work is needed to implement it into the system fully, but we have presented here a suggestion to the reader for advanced ways to normalise this assay.

## 4 Materials and Methods

The halo assay and cell culture were followed as per Laidlaw *et al*. [19]:

For the growth of yeast cultures, synthetic complete (SC) minimal medium (2% glucose and yeast nitrogen base supplemented with appropriate amino acids and bases) or rich yeast extract peptone dextrose (YPD) medium (2% glucose, 2% peptone and 1% yeast extract) were used. Plates were made using 2% agar, added before autoclaving. A shaker set to 225rpm at 30°C was used to grow cells overnight to early-mid log phase.

OD_600*nm*_ measurements were used to normalise lawn densities of strains grown to saturation overnight. Lawns were plated on methylene blue agar medium containing phosphate citrate buffer adjusted to pH A separate culture of the s*ki2*Δ*/ski2-2* diploid strain infected with the M28 virus was cultured to log phase (OD_600*nm*_ ≈ 1.0) before concentration to OD_600*nm*_=10, with 4 technical repeats per plate spotted on to the lawns. Yeast were grown for 48h at room temperature before growth was recorded.

Plates were imaged using an Epson Perfection 4990 scanner using the EPSON Scan software. For those wishing to perform their own experiments, we recommend using a scanner with a charge-coupled device (CCD) as opposed to a contact image sensor (CIS) as this gives better 3D contrast, essential when scanning thick agar plates.

## Code Availability

All code is made available at the following respository: https://github.com/PoL-biophysics/HaloUMIA static version of HaloUMI can be found via Zenodo [31].

## Data Availability

All images used to test, validate, and visualise are shown in training set folders via Zenodo [31]. The resulting figures are also included.

## Author Contributions

**Alex Pembery**: Conceptualisation, data curation, analysis, software, validation, visualisation, writing. **Hatwan Nadir**: Data curation, methodology. **Chris MacDonald**: Methodology, funding acquisition, supervision, writing. **Mark Leake**: Funding acquisition, supervision, writing

## Acknowledgements

This research was supported by a Sir Henry Dale Research Fellowship from the Wellcome Trust and the Royal Society 204636/Z/16/Z (CM) and by an EPSRC Fellowship EP/Y000501/1 (MCL).

## References

1. Hudzicki, J. Kirby-Bauer Disk Diffusion Susceptibility Test Protocol (American Society for Microbiology, 2009).

2. Schiller, H., Young, C., Schulze, S., Tripepi, M. & Pohlschroder, M. A Twist to the Kirby-Bauer Disk Diffusion Susceptibility Test: an Accessible Laboratory Experiment Comparing Haloferax volcanii and Escherichia coli Antibiotic Susceptibility to Highlight the Unique Cell Biology of Archaea. Journal of Microbiology & Biology Education 23, e00234–21. ISSN: 1935-7877 (2025).

3. Ong K. S. Screening of antibiotic sensitivity, antibacterial and enzymatic activities of microbes isolated from ex-tin mining lake. African Journal of Microbiology Research 5. ISSN: 19960808 (2011).

4. McCarty, T. P., Luethy, P. M., Baddley, J. W. & Pappas, P. G. Clinical utility of antifungal susceptibility testing. JAC-Antimicrobial Resistance 4, dlac067. ISSN: 2632-1823 (2022).

5. Yang, X. et al. Antimicrobial susceptibility testing of Enterobacteriaceae: determination of disk content and Kirby-Bauer breakpoint for ceftazidime/avibactam. BMC Microbiology 19, 240. ISSN: 1471-2180 (2019).

6. Balouiri, M., Sadiki, M. & Ibnsouda, S. K. Methods for in vitro evaluating antimicrobial activity: A review. Journal of Pharmaceutical Analysis 6, 71–79. ISSN: 2095-1779 (2016).

7. Barry, A. L., Coyle, M. B., Thornsberry, C., Gerlach, E. H. & Hawkinson, R. W. Methods of measuring zones of inhibition with the Bauer-Kirby disk susceptibility test. Journal of Clinical Microbiology 10, 885–889 (1979).

8. Petrova, M. & Petrov, P. A NEW METHOD FOR MANUAL MEASUREMENTS OF INHIBITION ZONES WITH THE BAUER-KIRBY DISK SUSCEPTIBILITY TEST in (2021).

9. Andrews, J. M., Boswell, F. J. & Wise, R. Evaluation of the Oxoid Aura image system for measuring zones of inhibition with the disc diffusion technique. Journal of Antimicrobial Chemotherapy 46, 535–540. ISSN: 0305-7453 (2000).

10. Le Page, S., Dubourg, G. & Rolain, J.-M. Evaluation of the Scan® 1200 as a rapid tool for reading antibiotic susceptibility testing by the disc diffusion technique. Journal of Antimicrobial Chemotherapy 71, 3424–3431. ISSN: 0305-7453 (2016).

11. Gullu, E., Bora, S. & Beynek, B. Exploiting Image Processing and Artificial Intelligence Techniques for the Determination of Antimicrobial Susceptibility. Applied Sciences 14. Number: 9, 3950. ISSN: 2076-3417 (2024).

12. Karayiğit, M., Erdem, O. & Güdücüoğlu, H. Artificial Intelligence Based Antibiotic Zone Measurement For Disk Diffusion in 2024 Signal Processing: Algorithms, Architectures, Arrangements, and Applications (SPA) 2024 Signal Processing: Algorithms, Architectures, Arrangements, and Applications (SPA) (2024), 48–53.

13. Kim, D., Lee, J. & Yoon, J. Accurate estimation of the inhibition zone of antibiotics based on laser speckle imaging and multiple random speckle illumination. Computers in Biology and Medicine 174, 108417. ISSN: 0010-4825 (2024).

14. Balmages, I. et al. Laser speckle imaging for visualization of hidden effects for early detection of antibacterial susceptibility in disc diffusion tests. Frontiers in Microbiology 14. ISSN: 1664-302X (2023).

15. Ramer, S. & Davis, R. A dominant truncation allele identifies a gene, STE20, that encodes a putative protein kinase necessary for mating in Saccharomyces cerevisiae. Proceedings of the National Academy of Sciences of the United States of America 90, 452–6 (1993).

16. Billerbeck, S., Walker, R. S. K. & Pretorius, I. S. Killer yeasts: expanding frontiers in the age of synthetic biology. Trends in Biotechnology 42, 1081–1096. ISSN: 0167-7799 (2024).

17. Becker, B. & Schmitt, M. J. Yeast Killer Toxin K28: Biology and Unique Strategy of Host Cell Intoxication and Killing. Toxins 9, 333. ISSN: 2072-6651 (2017).

18. Suzuki, Y., Schwartz, S. L., Mueller, N. C. & Schmitt, M. J. Cysteine residues in a yeast viral A/B toxin crucially control host cell killing via pH-triggered disulfide rearrangements. Molecular Biology of the Cell 28 (ed Linstedt, A.) 1123–1131. ISSN: 1059-1524, 1939-4586 (2017).

19. Laidlaw, K. M. E. et al. Killer toxin K28 resistance in yeast relies on COG complex-mediated trafficking of the defence factor Ktd1. Journal of Cell Science 138, jcs263897. ISSN: 0021-9533 (2025).

20. Prins, R. C. & Billerbeck, S. Protocol to study toxic, immune, and suicidal phenotypes of killer yeast. STAR Protocols 6. ISSN: 2666-1667 (2025).

21. Carroll, S. Y. et al. A Yeast Killer Toxin Screen Provides Insights into A/B Toxin Entry, Trafficking, and Killing Mechanisms. Developmental Cell 17, 552–560. ISSN: 15345807 (2009).

22. Costa, L. F. R. et al. Development of an automatic identification algorithm for antibiogram analysis. Computers in Biology and Medicine 67, 104–115. ISSN: 0010-4825 (2015).

23. Gerstein, A. C., Rosenberg, A., Hecht, I. & Berman, J. diskImageR: quantification of resistance and tolerance to antimicrobial drugs using disk diffusion assays. Microbiology 162, 1059–1068. ISSN: 1350-0872 (2016).

24. Bradski, G. The OpenCV Library. Dr. Dobb’s Journal of Software Tools (2000).

25. Virtanen, P. et al. SciPy 1.0: Fundamental Algorithms for Scientific Computing in Python. Nature Methods 17, 261–272 (2020).

26. Andreev, I. et al. Discovery of a rapidly evolving yeast defense factor, KTD1, against the secreted killer toxin K28. Proceedings of the National Academy of Sciences 120, e2217194120 (2023).

27. Gefen, O., Chekol, B., Strahilevitz, J. & Balaban, N. Q. TDtest: easy detection of bacterial tolerance and persistence in clinical isolates by a modified disk-diffusion assay. Scientific Reports 7, 41284. ISSN: 2045-2322 (2017).

28. Brook, I. Inoculum effect. Reviews of Infectious Diseases 11, 361–368. ISSN: 0162-0886 (1989).

29. Berg, H. C. Random Walks in Biology: New and Expanded Edition REV - Revised. ISBN: 978-0-691-00064-0 (Princeton University Press, 1993).

30. Carslaw, H. & Jaeger, J. Conduction of Heat in Solids ISBN: 978-0-19-853368-9 (Clarendon Press, 1959).

31. Pembery, A. HaloUMI Code, Figures, and Training Data (2026).

